# Highlighting the hidden: A tagging strategy for monitoring the association of GABARAP with microtubules in living cells

**DOI:** 10.1101/2023.11.22.568294

**Authors:** Alina Üffing, Lisa Gold, Thomas Gensch, Oliver H. Weiergräber, Silke Hoffmann, Dieter Willbold

**Affiliations:** Heinrich-Heine-Universität Düsseldorf, Mathematisch-Naturwissenschaftliche Fakultät, Institut für Physikalische Biologie, 40225 Düsseldorf, Germany; Forschungszentrum Jülich, Institut für Biologische Informationsprozesse: Strukturbiochemie (IBI-7), 52425 Jülich, Germany; Forschungszentrum Jülich, Institut für Biologische Informationsprozesse: Molekulare und Zelluläre Physiologie (IBI-1), 52425 Jülich, Germany

**Keywords:** ATG8, GABARAP, Live-cell imaging, Microtubules, Split-tandem, Tagging

## Abstract

GABARAP, like other ATG8 proteins, is a ubiquitin-like modifier and its C-terminal lipid conjugation enables association with cellular membranes. To prevent interference with the lipidation process, N-terminal fluorescent protein (FP) tagging strategies have become the standard for studying ATG8 localization and function in living cells, significantly contributing to our understanding of this protein family’s multifaceted roles.

However, recent findings have unveiled potential limitations of bulky N-terminal tags, particularly regarding ATG8 functionality and localization in specific contexts.

This study employed live cell imaging with particular emphasis on the GABARAP split-tandem construct, GABARAP(G116A)-mTagBFP2-GABARAP (G-*B*-G), which retains both a free N-terminus and a lipidation-competent C-terminus. Notably, our results revealed a robust association of G-*B*-G with the microtubule network in living cells which was not observed with N-terminal FP fusions of GABARAP, although early *in vitro* studies demonstrated an interaction of GABARAP and tubulin. Since we observed alteration of the microtubule network organization for G-*B*-G, this construct emerges as a valuable tool, which can help shedding light on potential roles of GABARAP in microtubule-associated processes that are integral to autophagy-related and -unrelated cellular transport.

## Introduction

The γ-aminobutyric acid type A receptor associated protein GABARAP belongs to the autophagy related protein 8 (ATG8) family, which in humans comprises a total of seven proteins. Human ATG8s are commonly split into two subfamilies, one containing the microtubule-associated protein 1 light chain 3 (MAP1LC3, hereafter referred to as LC3) isoforms LC3A, LC3B, LC3B2, LC3C, and one containing GABARAP, GABARAPL1, GABARAPL2. Like all ATG8 proteins, GABARAP consists of two N-terminal alpha-helices followed by a ubiquitin-like fold, and can similarly to ubiquitin be covalently conjugated to a substrate by an E1-E2-E3-like enzyme cascade (Fracchiolla et al., 2020; Komatsu et al., 2001; Lystad et al., 2019; Stangler et al., 2002; Tanida et al., 2002). Prior to substrate conjugation, ATG8s’ C-termini require processing by an ATG4 cysteine protease resulting in the exposure of a terminal glycine residue (e.g. G116 in GABARAP or G120 in LC3B). Deconjugation is also mediated by ATG4 proteases, making the process reversible (Fujita et al., 2008; Li et al., 2011; Tamargo-Gomez et al., 2021). Unlike ubiquitin, which is conjugated to amino groups in proteins, ATG8s usually exploit phosphatidylethanolamine (PE) or -serine (PS) as substrates, localizing them to membranes when lipidated (Ichimura et al., 2000). However, exceptions have recently been described for both types of modifiers (Agrotis et al., 2019; Carosi et al., 2021; Sakamaki et al., 2022).

To avoid conflict with the cellular lipidation process, N-terminal tagging strategies have been highly recommended for all ATG8s in the past (Klionsky, 2011), because C-terminal tagging would either result in loss of the tag or, in case of ATG4 processing-resistant ATG8 mutants like GABARAP(G116A), prevent membrane conjugation (Klionsky, 2011; Klionsky et al. 2021). Thus, N-terminal FP fusions have been widely used to study localization and function of ATG8s in living cells, and in conjunction with other techniques, have helped to elucidate the versatile roles of ATG8 proteins and shaped the current understanding of this multi-facetted protein family, which has been associated with a, still growing, plethora of autophagy related and unrelated functions (Heckmann et al., 2019; Leidal et al., 2020; Martinez et al., 2011; Solvik et al., 2022).

Interestingly, most of the underlying mechanisms have in common that they involve direct interactions of the ATG8s with other proteins. In the majority these interactions rely on a conserved motif in the interaction partner (LC3-interacting region, LIR; ATG8/GABARAP-interacting motif, AIM/GIM) and its corresponding docking site (often termed LDS for LIR-docking site) spanning two hydrophobic pockets on the ATG8 protein surface (Behrends et al., 2010; Wirth et al., 2019).

While it appears that many LIR/LDS-mediated ATG8-associated processes tolerate the presence of bulky, N-terminal tags, a growing body of evidence demonstrates that common ATG8 constructs, with bulky FP tags twice their size, can be functionally compromized and/or mislocalized. One example are studies on mitophagy, where N-terminal FP-tags were detrimental to GABARAP’s localization to mitochondria (Lazarou et al., 2015; Nguyen et al., 2016), while the smaller haemagglutinin-tag (HA-tag) did not inhibit mitochondrial targeting (Nguyen et al., 2016). The unique N-terminal region of ATG8 proteins distinguishes them from other ubiquitin-like proteins and its primary structure also varies remarkably among the different GABARAPs/LC3s. Notably, structural biology studies indicate that, contrary to the conserved and rather rigid ubiquitin-like cores, the N-termini of several human ATG8s, like the yeast Atg8 ancestor, exhibit high flexibility (Krichel et al., 2019; Schwarten et al., 2010; Stangler et al., 2002); (Möckel et al., 2019), a property that is likely to be altered in the presence of bulky tags. The N-terminal region of GABARAP, for instance, is known to adopt multiple conformations (Knight et al., 2002; Stangler et al., 2002) and has been described to be involved in its self-association (Coyle et al., 2002; Pacheco et al., 2010), in the regulation of its proteasomal degradation through MIB1-mediated ubiquitination of K13 and K23 (Joachim et al., 2017), and in its membrane association (Zhang et al., 2023). However, one of the first described functional features of the GABARAP N-terminal region was its tubulin and microtubule (MT) binding activity *in vitro*, which has also been documented for endogenous GABARAP in fixed cells by immunostaining (Wang & Olsen, 2000; Wang et al., 1999). Notably, FP-GABARAP usually shows a diffuse distribution pattern in cells under basal conditions, and can redistribute into punctate structures upon autophagy induction (Kabeya et al., 2004; Park et al., 2019; Tanida et al., 2006). To our knowledge, the description of a MT association of FP-GABARAP in living cells is still pending. Since systematic studies on ATG8-tagging strategies in general have received little attention in the past, we sought to fill this gap by employing a live cell imaging approach using cells expressing different FP arrangements of GABARAP. In contrast to the popular FP-GABARAP orientation, the presented work focuses on a so-called GABARAP split-tandem construct, GABARAP(G116A)-mTagBFP2-GABARAP (G-*B*-G). In G-*B*-G, an ATG4-resistant GABARAP with G116A substitution is joined to a central FP (mTagBFP2) followed by another GABARAP.

With this construct that combines a free GABARAP N-terminus with a lipidation competent C-terminus, we were able to shed first light on functions and localizations of GABARAP which otherwise might be hidden. Notably, we were able to observe a robust association of G-*B*-G with the MT network during live cell imaging. Given that MTs fulfill important roles in autophagy-related and -unrelated transport (Hilverling et al., 2022; Kimura et al., 2008; Labonte et al., 2014; Leil et al., 2004; Nakajima et al., 2012), our G-*B*-G construct might open new avenues to elucidating the involvement of GABARAP in these processes.

## Material & Methods

### DNA constructs

Plasmids used throughout this study are listed in table 1. All newly introduced vectors were generated by restriction-ligation cloning into pcDNA5/FRT/TO. Synthetic DNA fragments encoding GABARAP(G116A)-mTagBFP2-GABARAP (G-*B*-G), GABARAP(5X/G116A)-mCherry-GABARAP (G^5X^-*mCh*-G) and GABARAP(G116A)-mTagBFP2 (G-*B*) were obtained from Geneart and BioCat. All used constructs were sequence verified (Microsynth Seqlab). The respective plasmid DNAs have been deposited at the Addgene plasmid reposatory, where more detailed information for each construct can be accessed.

**Table 1.** List of constructs used in this work, together with short names, vector backbone, protein encoded, details (e.g. position-specific amino acid substitutions), and reference or source. Abbreviations: G: GABARAP; B: mTagBFP2; mCh: mCherry; Y: eYFP; *: G116A or G120A.

| Short names | Vector | Encoding | Details | Reference/Source |
| --- | --- | --- | --- | --- |
| *G-B-G* | pcDNA5/FR<br>T/TO | GABARAP(G116A)-mTagBFP2-GABARAP |  | Addgene # 212106 |
| *G<sup>5X</sup>-B-G* | pcDNA5/FR<br>T/TO | GABARAP(5X/G116A)-mTagBFP2-GABARAP | 5X: K2,<br>K13, R15,<br>K20, K23 to<br>A | Addgene # 212107 |
| *G-B-G\** | pcDNA5/FR<br>T/TO | GABARAP(G116A)-mTagBFP2-GABARAP(G116A) | lipidation-deficient | Addgene # 212108 |
| *B-G-G* | pcDNA5/FR<br>T/TO | mTagBFP2-GABARAP(G116A)-<br>GABARAP |  | Addgene # 212109 |
| *B-G* | pcDNA5/FR<br>T/TO | mTagBFP2-GABARAP |  | Addgene # 212110 |
| *Y-G* | pcDNA5/FR<br>T/TO | eYFP-GABARAP |  | Addgene # 212111 |
| *G-B* | pcDNA5/FR<br>T/TO | GABARAP(G116A)-mTagBFP2 |  | Addgene # 212112 |
| *G-mCh-G* | pcDNA5/FR<br>T/TO | GABARAP(G116A)-mCherry-GABARAP |  | Addgene # 212113 |
| mEos-tubulin | mEos3.2-C1 | mEos3.2-Tubulin-C-18 |  | Addgene # 57484 |

### Cell culture and transfection

Human hepatoma Huh-7.5 GABARAP SKO cells, HEK293 Flp in TRex GABARAP SKO cells (Dobner et al., 2020) and HeLa cells were cultured in Dulbecco’s Modified Eagle Medium (DMEM) high glucose (D5796, Sigma Aldrich), supplemented with 10% heat inactivated fetal bovine serum (F9665, Sigma) and 1% penicillin/streptomycin (P4333, Sigma Aldrich) at 37°C and 5% CO_2_. Cells were passaged at 80% confluency and routinely checked for mycoplasma contamination.

For transfection, cells were grown in 6-or 12-well plates to at least 70% confluency and transfected with 3 or 1.5 µg plasmid DNA per well, using 9 or 4.5 µl Lipofectamin 2000 (11668019, Thermo Scientific) respectively. Cells were seeded into fibronectin (F1141, Sigma) coated 35 mm glass-bottom IBIDI dishes 1.5-3 h post transfection and either imaged after 24-42 h or fixed for immunocytochemistry.

### Confocal Laser Scanning Microscopy

Cells were imaged using a LSM 710 confocal laser scanning system (Zeiss), operated with ZEN black 2009 software and a Plan-Apochromat 63x/1.40 Oil DIC M27 objective. For live-cell imaging at 37°C, a temperature-controlled microscopy stage was used. For staining of actin and microtubules, medium was supplemented with 500 nM SiR-tubulin (SC002, Spirochrome) or 1 µM SiR-actin (SC001, Spirochrome) and cells were incubated for 4 to 8 h prior to imaging. In the case of Huh-7.5 GABARAP SKO and HEK293 Flp in TRex GABARAP SKO, cells were additionally treated with 10 µM verapamil. The laser excitation wavelength and emission filters for the cells expressing fusion protein constructs were 405 nm and 410-509/530 nm for mTagBFP2, 514 nm and 519-621 nm for eYFP, 543 nm and 578-696 nm for mCherry and 633 nm and 638-759 nm for cells stained with SiR-probes.

### Colocalization colormap

To obtain a spatial representation of the colocalization of G-*B*-G with SiR-tubulin or SiR-actin we used the “colocalization colormap” Plugin for ImageJ (Gorlewicz et al., 2020), which is based on the Jaskolski algorithm (Jaskolski et al., 2005). This method creates a pseudo-color map of correlations between pairs of corresponding pixels in two input images. As input for the analysis we used smoothed images (3×3 mean filter) of G-*B*-G-expressing Huh-7.5 GABARAP SKO cells counter-stained either with SiR-tubulin or SiR-actin. Per analysed cell, five regions of interest (ROIs) of 40×40 pixels were split into their corresponding mTagBFP2- and SiR-channels. For each ROI the normalized mean deviation product (nMDP) was calculated for all corresponding pixels in the two channels, mathematically representing correlation between intensities of each pixel pair. The distribution of the calculated nMDP values was plotted, resulting in a colocalization colormap for each ROI. Such maps display the spatial correlation between the two fluorescent signals (mTagBFP2 and SiR), and because a jet colormap is implemented in the plugin by default, hot colours indicate colocalization and cold colours indicate separation. As a quantitative measure, the plugin also calculates the index of correlation (Icorr), which represents the fraction of positively colocalized pixels in each analyzed ROI. Finally, in order to compare the results, the distribution of Icorr values obtained for either G-*B*-G/SiR-tubulin or G-*B*-G/SiR-actin were plotted separately, and were subjected to statistical evaluation.

### Quantitative analysis of N/C, Nu/N and F/C ratios

To quantify the nucleocytoplasmic (N/C) ratio, five randomly ROIs (⌀ 10 px) were manually drawn in the nucleus as well as in the puncta-devoid cytoplasmic region of each cell using ImageJ. Mean intensities of nuclear and cytoplasmic ROIs from mTagBFP2 and eYFP channels were measured and divided. Likewise, mean intensities from 2-3 nucleoli ROIs were measured and divided by the mean nuclear intensity for both channels to determine the nucleolar to nucleoplasm (Nu/N) ratio of each cell. Filament to cytoplasm (F/C) ratios were determined by selecting 2-3 regions surrounding filaments using the polygon selection tool in ImageJ. For each filament ROI, a corresponding ROI of the same shape and size was placed in a nearby cytoplasmic region without filaments. Mean intensities were measured and F/C ratios calculated per ROI pair and cell. In case of co-transfected cells, this was done for both the mTagBFP2 and the eYFP channel. In case of single construct transfection and co-staining with SiR-tubulin, ROI pairs were selected according to most intense signals in the SiR channel and mean intensities were measured from the mTagBFP2 channel. N/C, Nu/N and F/C ratios were plotted and subjected to statistical evaluation.

### Microtubule organization analysis

For analysis of the microtubule cytoskeleton, images showing a G-*B*-G expressing and an un-transfected control cell (co-)stained with SiR-tubulin were selected. The areas of both individual cells were manually traced according to the transmission image and saved as ROIs. SiR channels were skeletonized using the ImageJ LPX filter2D plugin (filter = lineFilters, linemode = lineExtract, giwsiter = 5, mdnmsLen = 15, pickup = otsu, shaveLen = 5, delLen = 5; (Ueda et al., 2010). Afterwards, the skeleton image type was set to 8 bit, duplicated, ROIs selected and background set to black (0,0,0) to obtain a skeletonized microtubule image per cell (ROI). Subsequently, the skeleton of each cell was analysed using ImageJ (Analyze > Skeleton) to tag endpoint, junction and slab pixels (the pixels between junctions and endpoints). Branch length was plotted for each cell and junctions were divided by total length to obtain a junctions per µm of branch (j/µm) value for each cell.

Additionally, cytoskeleton bundling parameters (Skewness and coefficient of variation [CV]) were obtained from the skeleton images of each cell according to an ImageJ macro published by (Higaki et al., 2020), modified for analysis of single plane images.

### Prediction of complex stuctures

Potential modes of interaction of human GABARAP or its mTagBFP2 tandem constructs with microtubules were investigated *in silico* with AlphaFold2 (Evans et al., 2022 preprint), using the ColabFold implementation (Mirdita et al., 2022). Complexes of GABARAP and a TBA1A-TBB5 (Uniprot accession Q71U36, P07437) dimer were predicted using a local installation (github.com/YoshitakaMo/localcolabfold) running on a Linux workstation equipped with an nVidia GPU, while data for the larger complexes of G-*B*-G or *B*-G-G with a tetrameric tubulin chain (TBA1A-TBB5-TBA1A-TBB5) were uploaded to the COSMIC^2^ platform (Cianfrocco et al., 2017) for processing by ColabFold. For each subject, five models were generated with inclusion of template information, using multiple sequence alignment mode *mmseqs2_uniref_env*, model type *alphafold2_multimer_v3*, a maximum of 20 recycle steps, and recycling controlled by *recycle_early_stop_tolerance:0.5* and *stop_at_score:90* for the local and remote installations, respectively. All models were subjected to relaxation using Amber as implemented in the ColabFold pipeline. In general, the 1^st^ rank model, according to predicted lDDT statistics, is used for further evaluation; the complete list of models along with predicted lDDT and PAE plots is shown as supplemental information (Fig. S5).

### Statistical analysis

Mean N/C and Nu/N values per cell as well as resulting statistics from paired t-tests were plotted using GraphPad Prism version 9. Mean F/C values per cell were analyzed by paired t-test in case of co-transfected cells and one-way Anova with Tukey’s multiple comparison test for single transfections with different constructs using GraphPad Prism version 9, and mean values as well as individual values were plotted using SuperPlot (https://huygens.science.uva.nl/SuperPlotsOfData/). Mean Icorr values per cell were analyzed by unpaired t-test, and individual values as well as means were plotted using Graphpad Prism 9 and SuperPlot, respectively. Branch length was analyzed for each cell pair using Mann-Whitney test, and individual values were plotted with Graphpad Prism 9.

## Results

### G-*B*-G exhibits subcellular localization distinct from *Y*-G

Due to Atg8-like proteins being C-terminally conjugated to lipids, their N-terminal fusions with fluorescent proteins (FP-ATG8s), such as the yellow fluorescent protein-tagged GABARAP (*Y*-G) used in this study, have a long tradition in the study of their function in living cells. Autophagy induction typically triggers FP-ATG8s to localize to punctate structures, interpreted as their lipidated forms associated with autophagic membranes. Under basal conditions, being the focus here, FP-ATG8s are known for their diffuse cytoplasmic and nuclear distribution - commonly interpreted as their free, unlipidated forms. To restrict GABARAP functionalities to the transfected constructs, excluding any contribution from an endogenous GABARAP background, Huh-7.5 GABARAP knockout cells, hereafter abbreviated as Huh^KO^ cells, were used throughout this study unless otherwise stated. In order to be able to directly compare the behavior of the novel split tandem construct G-*B*-G with that of *Y*-G under basal conditions, cells were co-transfected with the corresponding expression plasmids (Table 1). Overexpressed *Y*-G showed, as expected, a diffuse cytoplasmic distribution with a clear enrichment in the nucleoplasm, however, co-expressed G-*B*-G appeared to accumulate on a variety of intracellular structures, resulting in a more distinct staining pattern (Figure 1A). Even though the mean nucleocytoplasmic (N/C) ratio of *Y*-G (1.69 ± 0.34) was significantly higher than for G-*B*-G (1.03 ± 0.45), the latter showed visible enrichment at diverse nuclear subcompartements including the nucleolus, with a higher mean nucleoli-to-nucleoplasm (Nu/N) ratio (2.27 ± 0.49) compared to *Y*-G (1.09 ± 0.11; Figure 1B-D).

**Figure 1.**
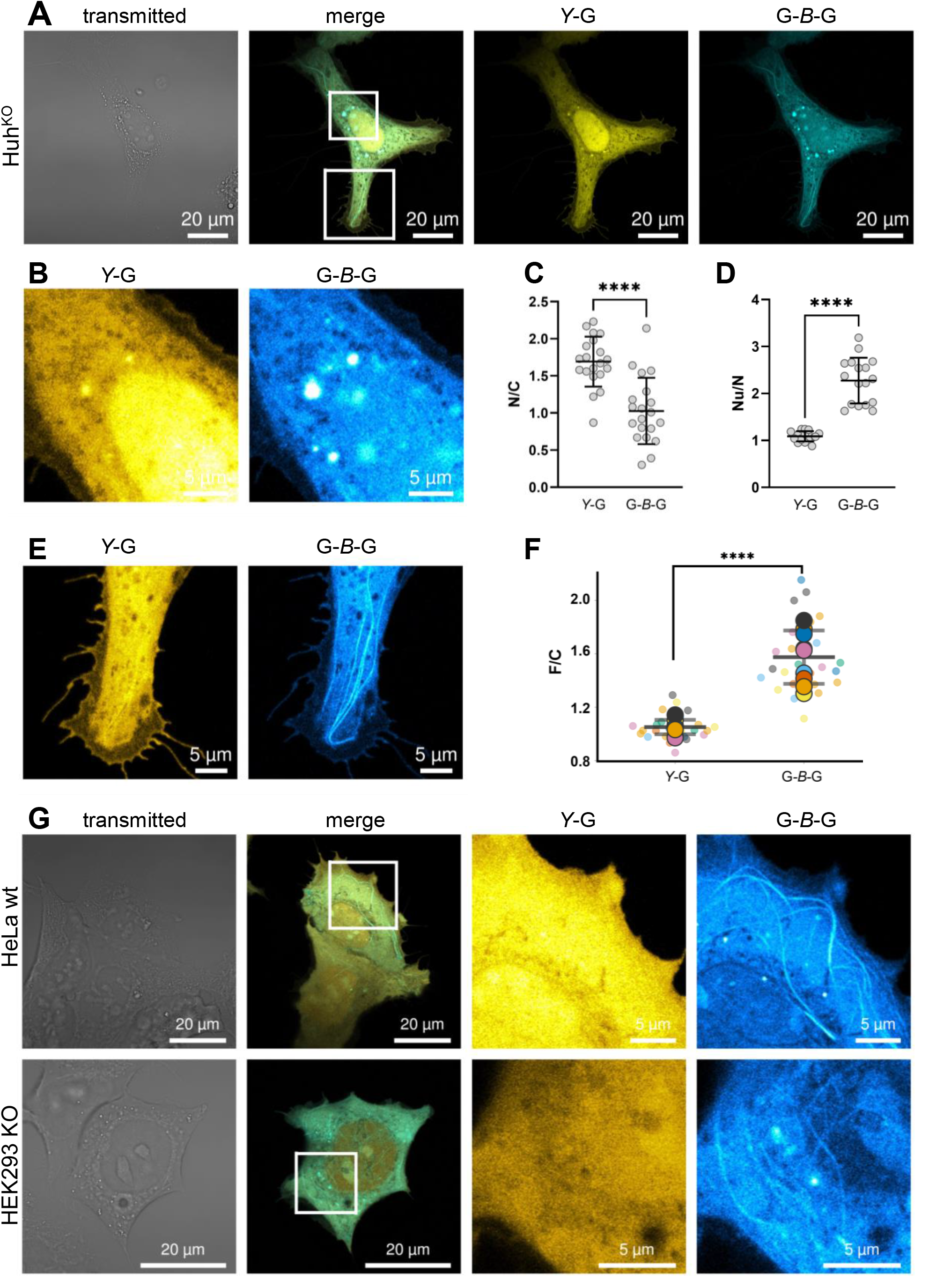
Distinct subcellular distribution of *Y*-G and G-*B*-G. (**A**) Exemplary image of a Huh7.5 GABARAP KO cell expressing G-*B*-G and *Y*-G. Distinct signals are visible at different subcellular locations including (**B**) nucleus and nucleoli and (**E**) filaments. Quantification of (**C**) nucleocytoplasmic ratio (N/C), Mean: 1.03 ± 0.45 for G-*B*-G and 1.69 ± 0.34 for *Y*-G; (**D**) Nucleoli to nucleoplasm (Nu/N) ratio, Mean: 2.27 ± 0.49 for G-*B*-G and 1.09 ± 0.11 for *Y*-G and (**F**) filament to cytoplasm ratios (F/C), Mean: 1.57 ± 0.2 for G-*B*-G and 1.05 ± 0.05 for *Y*-G measured by mean fluorescence intensity. **** P < 0.0001, paired t-test. Values are represented as means ± SD (n= 20 (N/C), 17 (S/N), 9 (F/C) from two independent experiments). See Fig. S1 for a detailed description of the quantification procedure applied. All cells and ROIs that went into the quantification can be reviewed on BioImageArchive. (**G**) In HEK293 GABARAP SKO and HeLa wildtype cells, distinct features can also be observed for overexpressed G-*B*-G compared to *Y*-G.

Most strikingly, G-*B*-G was frequently enriched at filamental structures while this was barely visible for *Y*-G in the respective cells (Figure 1E). The mean filament-to-cytoplasm ratio (F/C) for *Y*-G (1.05 ± 0.05) represents an absence of enrichment, in contrast to the increased F/C ratio for G-*B*-G (1.57 ± 0.2, Figure 1F). To assess whether the discrepancies in filament association observed between G-*B*-G and *Y*-G arose from the different FPs used, Huh^KO^ cells were transfected with a construct expressing N-terminally mTagBFP2-tagged GABARAP. However, like *Y*-G, *B*-G did not show the obvious filament association seen for G-*B*-G. Likewise, cells expressing a C-terminally tagged, lipidation-deficient GABARAP with a free and unaltered N-terminus, G-*B*, also failed to show any robust filament association, suggesting that the mere presence of an unmodified N-terminus is not sufficient to establish filament association of an FP-linked GABARAP or at least fails to achieve the degree of association sufficient for a robust microscopic detection. Interestingly, N-terminal tagging of two consecutive GABARAPs as in *B*-G-G resulted in a similar association with filaments as demonstrated for G-*B*-G (Fig. S1B-F). These observations suggest that combining two GABARAP moieties in a single polypeptide, e.g. flanking a central mTagBFP2, may be a promising way to microscopically study GABARAP’s filament association-related activities in living cells.

Since we observed pronounced G-*B*-G filament association particularly frequently in Huh^KO^ cells, this line was used for all further experiments. However, this phenomenon and the other distinct features of G-*B*-G and *Y*-G described above were also evident in other lines as illustrated for HEK293 cells in a GABARAP single-KO background, and for wildtype HeLa cells (Figure 1G).

### G-*B*-G-decorated filamental structures correspond to microtubules

To investigate the identity of the G-*B*-G enriched filamental structures, Huh^KO^ cells expressing G-*B*-G were co-stained with SiR-tubulin and SiR-actin. Representative images showed a broad match of the fluorescence signals for G-*B*-G with SiR-tubulin, additionally confirmed through mEos-tubulin colocalization (Fig. S2A, D), whereas G-*B*-G and SiR-actin signals overlapped poorly, at best appearing with a parallel offset or crossing each other (Figure 2A-B, Fig. S2B). Spatial correlation of G-*B*-G with SiR-tubulin and SiR-actin signals was quantitated as outlined in the Methods section; Icorr values determined for G-*B*-G/tubulin (0.68 ± 0.02) were significantly higher than those for G-*B*-G/actin (0.29 ± 0.09), confirming the abovementioned observation (Figure 2C-D). In most cases the enrichment of G-*B*-G along MTs was uniform, but in some cells G-*B*-G positive puncta, possibly G-*B*-G decorated transport vesicles, were observed in proximity to MTs (Fig. S2C, D), raising the idea of a connection between vesicle-associated GABARAP and MTs.

**Figure 2.**
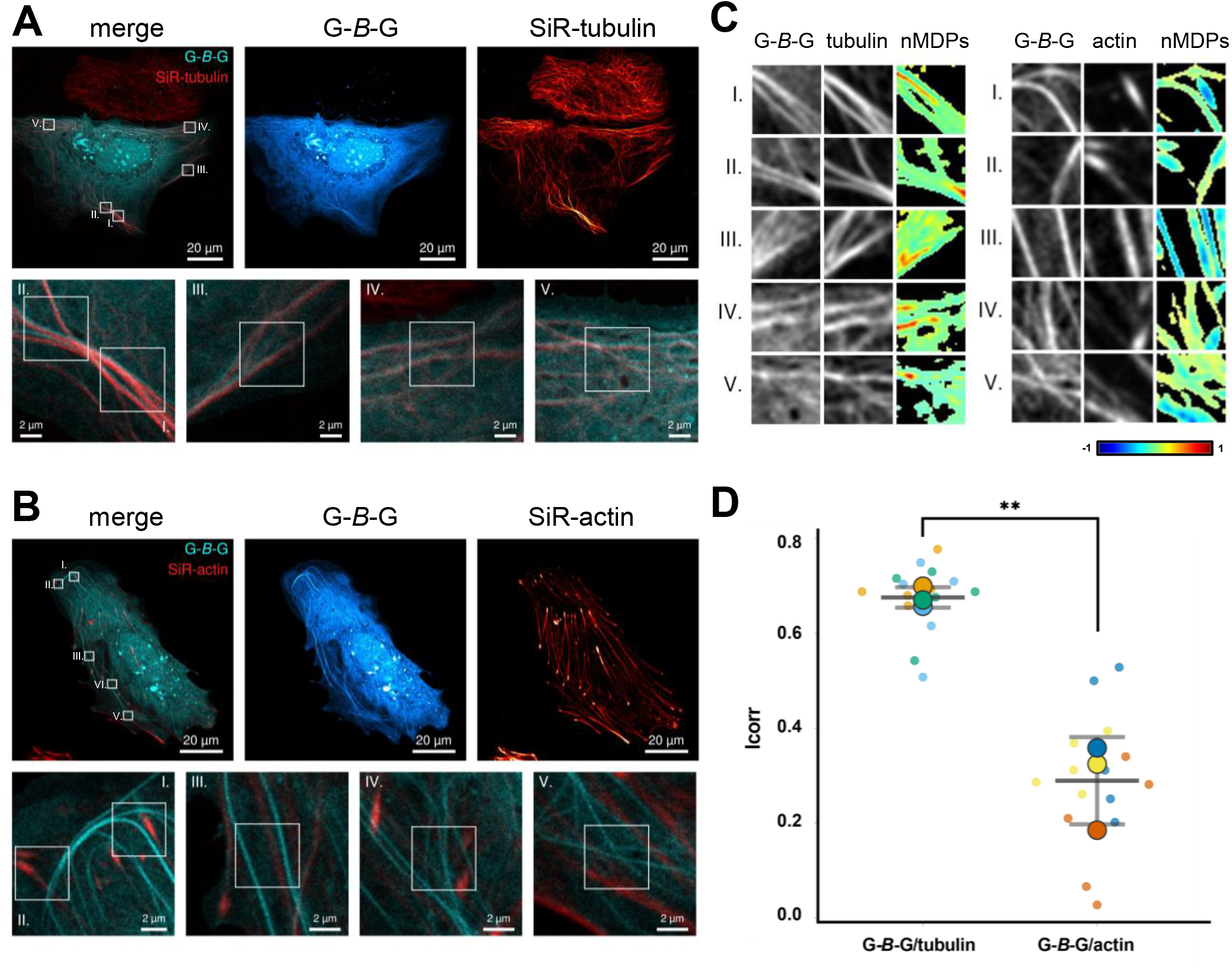
Filamental G-*B*-G structures correlate with SiR-tubulin signal. Representative live-cell images of G-*B*-G expressing Huh^KO^ cells stained with (**A**) SiR-tubulin or (**B**) SiR-actin. Bottom panels show magnifications around five selected ROIs per cell. (**C**) Grey-scale images of the ROIs depicted in (**A**) and (**B**) highlighting the distribution pattern of G-*B*-G and tubulin (left) or actin (right). The corresponding colocalization colormaps (nMDPs, 40×40 pixel) show the spatial correlation between the two fluorescent signals (mTagBFP2 and SiR), with hot colours indicating colocalization and cold colours indicating separation (n=3; refer to Fig. S3 for replicates). (**D**) Graphical representation of the corresponding Icorr-values representing the fraction of positively correlated pixels for G-*B*-G and tubulin or actin. Values are plotted both for each ROI (small dots; Icorrs of the same cell have the same colour) and as mean Icorr for all ROIs of a single cell (large dots). From the latter the overall Icorr mean and SD for G-*B*-G/tubulin (0.68 ± 0.02) and G-*B*-G/actin (0.29 ± 0.09) were calculated. ** P=0.0021; unpaired t-test.

### N-terminal residues support enrichment of G-*B*-G at microtubules while lipidation capability is dispensable

Having shown that G-*B*-G associates preferentially with MTs, we then asked which regions in GABARAP, or in G-*B*-G, might be involved. For this purpose, Huh^KO^ cells expressing G^5X^-*B*-G with a total of five alanine substitutions of N-terminal basic residues (K2A, K13A, R15A, K20A, K23A) in the first GABARAP moiety of the split tandem construct were investigated. These residues have been suggested to be part of GABARAP’s tubulin binding motif in former *in vitro* studies. Indeed, compared to control cells expressing G-*B-*G, G^5X^-*B*-G-expressing cells mostly showed little to no enrichment at MTs, which were visualized by co-staining with SiR-tubulin (Figure 3A, B & Fig. S3A, B). Additionally, Huh^KO^ cells expressing G-*B*-G* and co-stained with SiR-tubulin were imaged, permitting investigation of whether lipidation is required for MT association, as the corresponding G-*B*-G* fusion protein is devoid of C-terminal glycine residues suitable for lipidation in both GABARAPs. Interestingly, lipidation deficiency did not appear to inhibit enrichment of G-*B*-G* at MTs, as cells expressing this construct presented themselves indistinguishable from cells expressing the unmodified G-*B*-G (Figure 3A, C & Fig. S3C).

**Figure 3.**
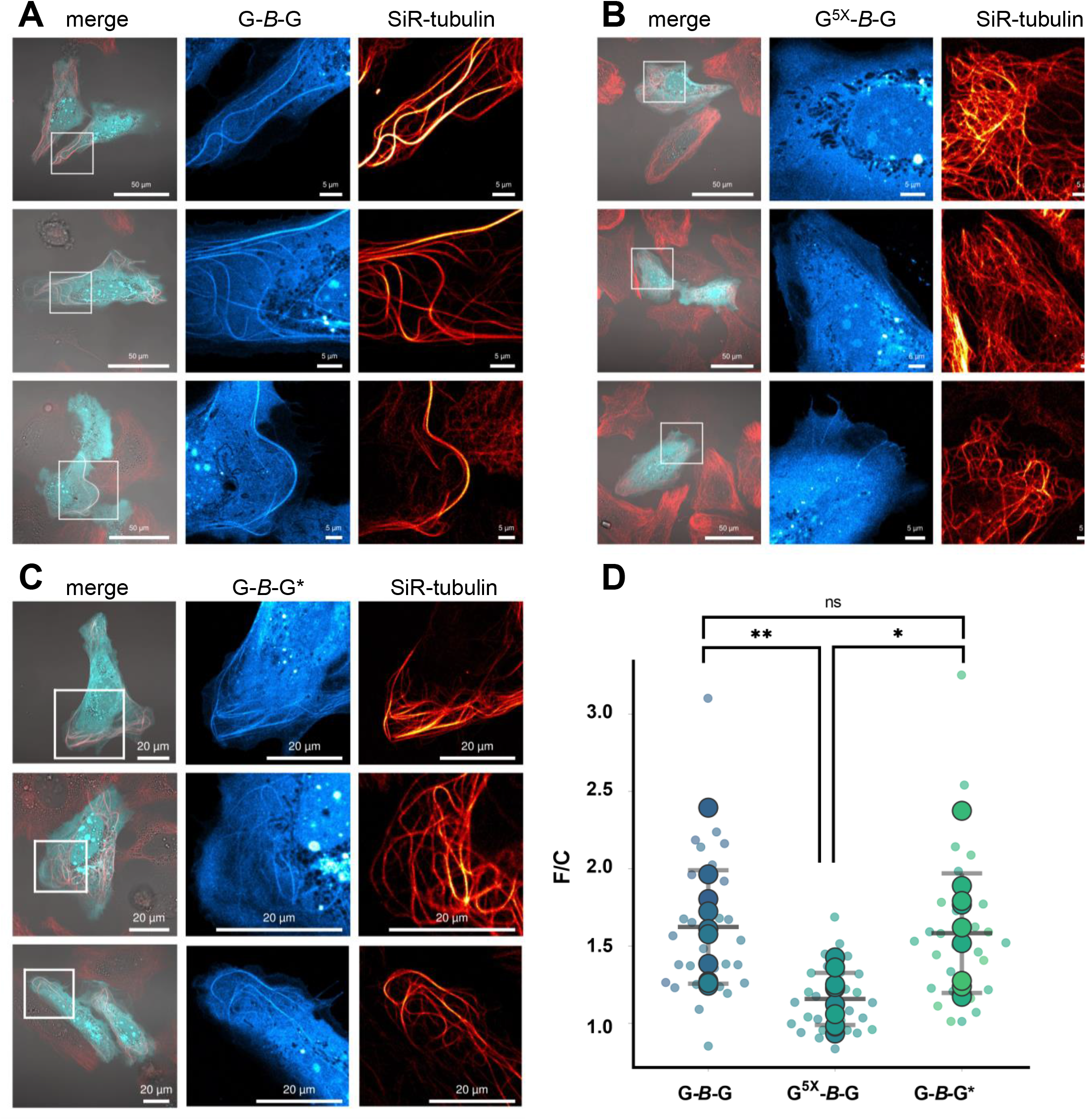
Reduced filament/microtubule association in the absence of conserved, positively charged residues in the tubulin-binding region. Exemplary image of a Huh^KO^ cells stained with SiR-tubulin expressing (**A**) G-*B*-G, (**B**) G^5X^-*B*-G and (**C**) G-*B*-G* (**D**) Quantification of filament to cytoplasm ratios (F/C) measured by mean mTagBFP2 fluorescence intensity at respective ROIs with strong SiR-tubulin signal (3/cell, small dots, mean per cell: large dots). Please refer to Fig. S3 for images of all cells that went into this quantification. Additionally, all cells and ROIs can be reviewed on BioImageArchive. One way Anova with Tukey’s multiple comparison test. ** P=0.0089, * P=0.0174 ns= non significant. Values are represented as means ± SD (n=10 cells) for each genotype resulting in 1.62 ± 0.37, 1.16 ± 0.168 and 1.68 ± 0.39 for G-B-G, G^5X^-*B*-G and G-*B*-G*, respectively.

These observations were again substantiated by quantitative analysis. While mean F/C ratios of G-*B*-G and G-*B*-G* expressing cells showed no significant difference (1.62 ± 0.37 and 1.68 ± 0.39), enrichment at filaments compared to adjacent cytoplasm was significantly lower (1.16 ± 0.168) for G^5X^-B-G (Figure 3D). Certainly, using different fluorophores and comparing the different constructs upon co-expression in the same cells, as shown for *Y*-G and G-*B*-G in Fig. 1, would have been the straightforward method to address the questions described above. Indeed, we intended to follow this approach, however, to our surprise, a split-tandem construct utilizing mCherry (mCh) as FP, G-*mCh*-G, did not show MT association comparable to G-*B*-G (Fig. S3D, E). As full-length translation and a good expression level of G-*mCh*-G, just as for any other tandem construct, was confirmed (Fig. S3F), the question of why MT association of GABARAP was less visible with G-*mCh*-G remains unanswered.

### G-*B*-G expression can alter microtubule network organization

During investigation of G-*B*-G expressing cells, it became apparent that G-*B*-G decorated MTs often appeared strikingly long and curved, with a tendency to even build loops. To analyze this prominent alteration of the MT network, Huh^KO^ cells expressing G-*B*-G were co-stained with SiR-tubulin. Pairs of untransfected cells and cells strongly expressing G-*B*-G were selected and analyzed regarding their MT network organization; we observed clear differences in the network throughout the analyzed cells, with G-*B*-G expressing cells presenting prominently long and curved MTs (Figure 4A). Skeletonization of filamental structures in the SiR channel and subsequent determination of branch length, defined as distance between endpoint and/or junction pixels, showed significantly longer branches for all analyzed G-*B*-G expressing cells (extending up to 50 µm) compared to control cells (Figure 4B-D, Fig. S4A). Additionally, cytoskeleton bundling parameters, namely coefficient of variance (CV) and skewness, were examined for pairs of G-*B*-G expressing and control cells with similar mean intensities in the SiR-channel (less than 1.6-fold difference, Image 1,2,5). Higher CV and Skewness values for G-*B*-G expressing cells compared to control cells supported the frequent visual observation of high SiR-tubulin intensities associated with pronounced, long MTs (Figure 4E-G, Fig. S4B). It is important to note that the altered MT network organization in G-*B*-G expressing cells was not only observable with SiR-tubulin staining, which is known to stabilize MTs, but also in fixed cells stained with an antibody against β-tubulin (Fig. S4C).

**Figure 4.**
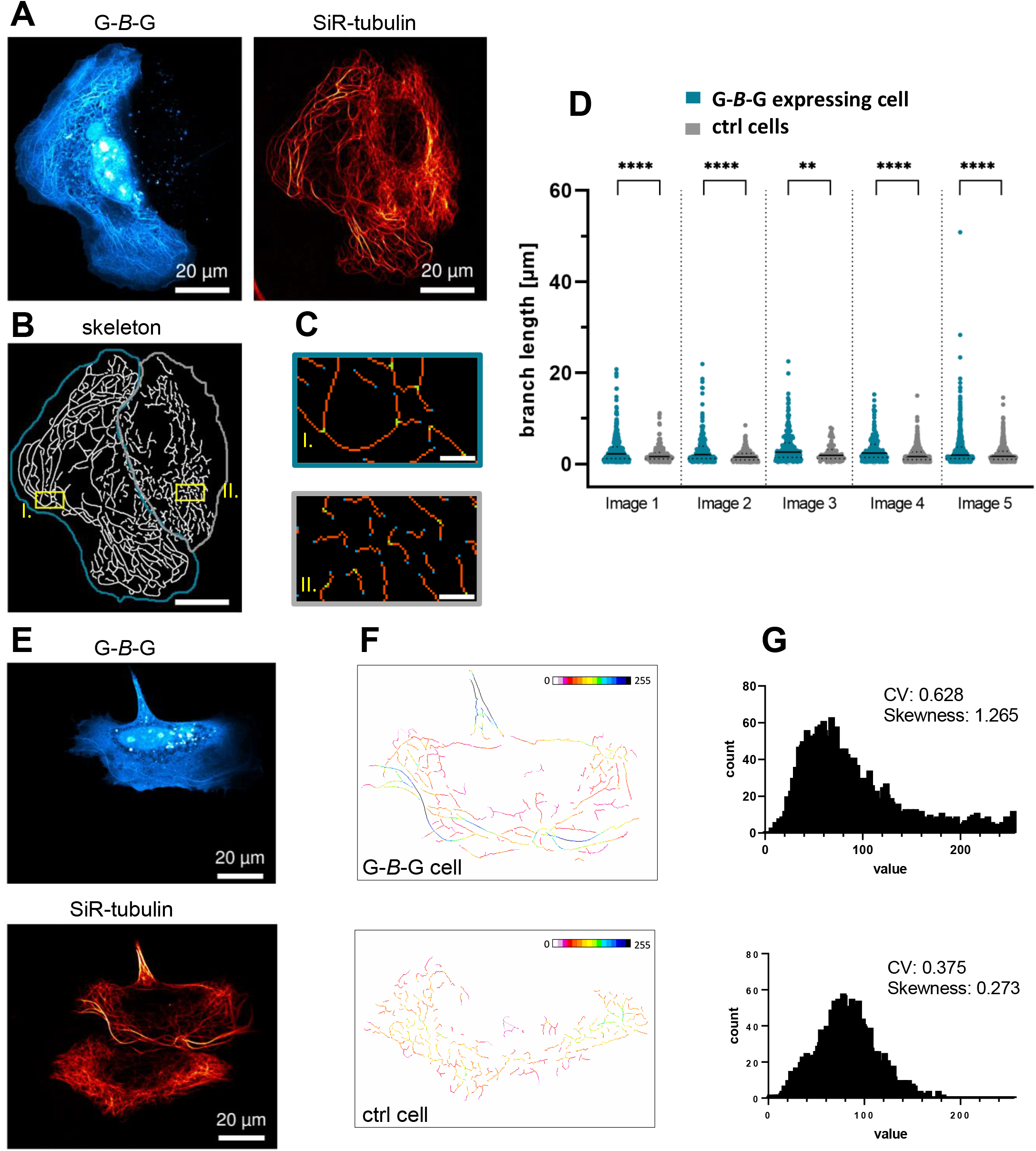
Altered microtubule structure/organization in G-*B*-G expressing Huh^KO^ cells. (**A**) Exemplary G-*B*-G expressing cell and control cell, both stained with SiR-tubulin (Image 1). (**B**) Skeleton of SiR-tubulin stain in Image 1. The G-*B*-G expressing cell is marked in blue, the control cell in gray. Skeleton was thresholded and dilated for visualization purposes. Exemplary regions are marked in yellow. (**C**) Zoomed in ROIs from skeletonized image after tagging of endpoint (blue), junction (green) and slab (orange) pixels for each cell. (**D**) Quantification of branch length from 5 images (see Fig. S7A) each displaying a G-*B*-G expressing and control cell. **** P < 0.0001, ** P=0.0021, Mann-Whitney test. Individual values are represented including medians and quartiles (**E**). Exemplary G-*B*-G expressing cell and control cell stained with SiR-tubulin (Image 2). (**F**) Skeletonized cells colored according to intensity of tubulin staining. (**G**) Histograms corresponding to skeletonized cells in F. Cytoskeleton bundling indicators (CV and Skewness) are shown for each cell. For two more examples refer to Fig. S4B.

### AlphaFold predictions suggest charge-mediated GABARAP enrichment at microtubules rather than a distinct binding mode

The presence of two GABARAP molecules in the fusion protein construct G-*B*-G appeared to be one critical factor for MT association. Using ColabFold to predict the structural arrangement of G-*B*-G with a protofilament composed of two tubulin dimers (TBA1A-TBB5), we sought to address the question whether this interaction is sterically plausible. First, orientation of the two GABARAP moieties towards the tubulin chain and of mTagBFP2 away from it was consistently observed, with the GABARAPs typically aligning with the tubulin protofilament in such a way that one protomer is skipped (Figure 5A, Fig. S5A). Despite significant variance among individual models regarding details of the predicted contacts, GABARAP was frequently suggested to associate with the negatively charged C-terminal tail present in both α- and β-tubulin. Due to the inherent disorder of this segment, GABARAP is unlikely to adopt a well-defined orientation relative to the globular tubulin core, as reflected by high PAE scores for intermolecular pairs of residues. With due caution owing to the complexity of the system, these predictions support the idea of electrostatic interactions between G-*B*-G and microtubules. Predictions of untagged GABARAP with a TBA1A-TBB5 dimer also hint at an attraction of GABARAP toward the exposed C-terminal tails of the tubulin monomers (Figure 5B. Fig. S5B). In accordance with the observation in cells, where *B*-G-G showed enrichment at MTs comparable to G-*B*-G, ColabFold predicted a similar charge-dominated tubulin interaction of GABARAP moieties, despite their shorter separation in *B*-G-G (Fig. S5C). However, predictions with untagged GABARAP and tetrameric tubulin did not yield consistent results, possibly indicating a lower interaction propensity. Notably, these calculations do not include the various post-translational modifications described for both GABARAP and tubulin and only represent tubulin dimers and tetramers of specific isotypes, TBB5 and TBA1A. However, both post-tranlational modifications and isotypes are important for structure and functions of MTs, as well as for interactions with other proteins, possibly including the proposed interaction with GABARAP.

**Figure 5.**
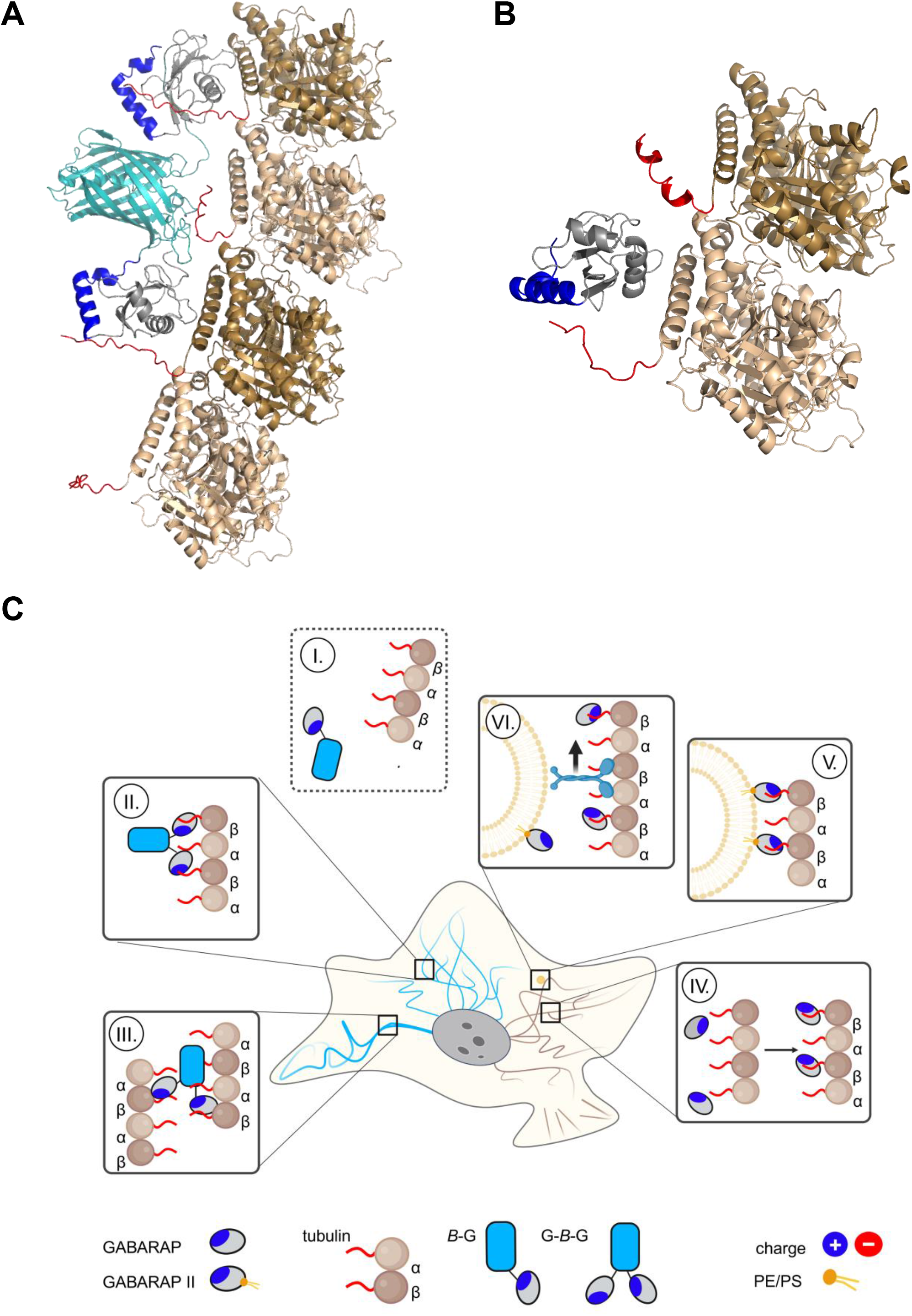
Predictions support a GABARAP-tubulin interaction, although FP and valency of GABARAP likely influence the biological outcome. Complex models of tubulin oligomers with G-*B*-G (**A**) and GABARAP (**B)**. This studýs observations along with their possible biological consequences are summarized schematically in (**C**), with more details being explained in the main text.

## Discussion

N-terminal FP tagging of ATG8 proteins has been a long-standing method for fast and easy monitoring of autophagy-related and -unrelated processes (Kabeya et al., 2000; Leil et al., 2004). Due to concerns about this method, there have been increasing efforts to develop further strategies for live-cell visualization of these proteins (Klionsky et al., 2021). Different sensors, either for individual ATG8 proteins (Stolz et al., 2017) or membrane associated ATG8s (Lee et al., 2017) have been generated with the aim to visualize endogenous ATG8 proteins. Both sensors and FP tags could influence ATG8s functions and localizations through occupation or alteration of binding sites. In line with this, a recent study using overexpressed tag-less GABARAP identified a *cis*-membrane binding activity of the N-terminal region of GABARAP as a prerequisite for autophagosome membrane expansion (Zhang et al., 2023).

Here, we introduced G-*B*-G, a GABARAP split-tandem construct, as a tool for live-cell imaging, which albeit possessing a large FP tag, displays a free N- and C-terminus and could therefore help to broaden the knowledge on localization and function of GABARAP.

In a majority of cells, we observed differences in the nucleocytoplasmic distribution between G-*B*-G and the conventionally labeled *Y*-G (expressed simultaneously or separately) with overall stronger accumulation of *Y*-G in the nucleoplasm. Pronounced nuclear localization has been interpreted as an artefact in some settings (Klionsky et al., 2021), however, ATG8 proteins have also been reported to interact with nuclear proteins including lamin B1, where N-terminally located residues were identified as important for the interaction (Dou et al., 2015), and TP53INP2, which in response to starvation shuttles nuclear deacetylated LC3B to the cytoplasm (Huang et al., 2015). Reported sensitivity to a deacetylase inhibitor suggests a similar mechanism for GABARAP (Baeken et al., 2020). Within nuclear compartments different accumulation behaviors have been described for LC3s upon immunodetection in diverse cancer cells lines (Koukourakis et al., 2015), and rapid nucleolus-nucleoplasm shuttling has been demonstrated for Venus-LC3B (Kraft et al., 2016). Interestingly, G-*B*-G was found to be strikingly enriched in nucleoli, whereas *Y*-G was not. This may indicate a previously overlooked and yet to be defined retention or attraction mechanism for GABARAP in this nuclear subcompartment, possibly supported by the presence of an available N-terminus in G-*B*-G.

In our view, the most striking feature of G-*B*-G is its ability to accumulate on filamentous structures, namely MTs, in the cytoplasm, a behavior hitherto undocumented for FP-tagged GABARAP/ATG8s. Interestingly, association of GABARAP with MTs has in fact been reported in fixed CHO cells more than 20 years ago (Wang & Olsen, 2000). One might wonder why this observation, obtained by immunofluorescence, has so far remained the only evidence for a possible intracellular GABARAP-MT association. Since the polyclonal anti-GABARAP antibody used at that time is no longer available, the original results cannot be reproduced. It is quite conceivable that the GABARAP epitope recognized by the antibody could be decisive for the result, and notably, using the monoclonal anti-GABARAP antibody 8H5 (Simons et al., 2019), we never noticed any obvious MT association of GABARAP, presumably because the N-terminally located epitope for this antibody overlaps with the proposed tubulin-binding region. Further possible explanations include too low GABARAP protein levels, an unfavorable cell state, and of course also an adverse effect of the tagging strategies used.

Contrary to the sparse data reported within cells, there is abundant evidence for a robust interaction of GABARAP both with tubulin and assembled MTs *in vitro*. In 2000, Wang and Olsen demonstrated association of GST-GABARAP with purified tubulin but not actin, and further showed interaction of heterologously expressed GABARAP with *in vitro* assembled MTs, supporting our observation that lipidation is not a prerequisite for G-*B*-G enrichment at MTs in cells, and narrowed down the responsible region to the first 35 residues of GABARAP (Wang & Olsen, 2000). Together with other results these data led to the suggestion that the positively charged residues in the N-terminal region of GABARAP promote its association with MTs through ionic interactions with the negatively charged C-terminal tails of the tubulin monomers (Coyle et al., 2002; Wang & Olsen, 2000).

Our results indicate that positively charged residues within the GABARAP N-terminus are crucial for a robust MT association also in the intracellular context, as we observed a significantly reduced MT association for G^5x^-*B*-G, which lacks the basic residues K2, K13, R15, K20 and K23 in the first GABARAP of the split tandem. It is known that GABARAP can promote MT polymerization with faster kinetics than the MT-stabilizing drug taxol *in vitro* and that the first 22 N-terminal residues are sufficient for this purpose (Coyle et al., 2002; Wang et al., 1999). Coyle et al. (2002) already suspected an MT-bundling activity of GABARAP based on their *in vitro* turbidity experiments, as the addition of GABARAP to single MTs, preassembled by taxol (Yoon & Oakley, 1995), led to a further strong increase in turbidity. Consistent with these results, a rapid increase in turbidity was reported after addition of the closely related GABARAP paralog GABARAPL1 to MgCl_2_ preassembled single MTs (Mansuy et al., 2004). Shielding of the negatively charged C-terminal tail of tubulin by positively charged patches on MT-associated proteins, as described for the cytoskeleton-associated protein glycine-rich (CAP-Gly) domain of p150^glued^ (Wang et al., 2014), offers one possible mechanism for these *in vitro* observations, since the basic N-termini of GABARAP and GABARAPL1 likewise could neutralize the repulsive negative surface charge of MTs. Interestingly, we repeatedly observed an altered MT network organization including cytoskeleton bundling in G-*B*-G expressing cells, indicating that this GABARAP activity may indeed have significance *in vivo*. Since the G-*B*-G construct offers two tubulin binding sites it also could bridge individual MTs to form bundles. The bundling promoting properties of the MT associated protein tau have been suggested to be based on a similar mechanism, namely the dimerization of MT bound tau (Feinstein et al., 2016). Remarkably, Nymann-Andersen et al. (2002) already discussed MT binding of GABARAP dimers after identification of the LDS as the self-association domain at physiological salt concentrations *in vitro*. How the observed MT network alterations are facilitated in detail, and to what extent these effects observed with the split tandem construct can be translated to endogenous levels of GABARAP is subject to speculation and requires further investigation.

Compared to the relatively controlled conditions *in vitro,* factors influencing MT dynamics and stability in living cells are much more complex. Stability determining factors include the prevalence of tubulin isotypes (Fu et al., 2023), different post-translational modifications (Bonnet et al., 2001; Eshun-Wilson et al., 2019; Konno et al., 2016; Portran et al., 2017; Tas et al., 2017), and the presence or absence of diverse MT associated proteins (MAPs, Cuveillier et al., 2020; Shigematsu et al., 2018), which often regulate transport along MTs. Tau is one example of a structural MAP that stabilizes and bundles MTs but inhibits kinesins and, though to a lesser degree, dyneins (Dixit et al., 2008; Kanai et al., 1992). The motor protein kinesin-1 itself has been shown to preferentially bind to more stable acetylated MTs in the perinuclear region, while kinesin-3 transports cargo on more peripheral, tyrosinated and thus more dynamic MTs (Guardia et al., 2016). Neuronal transport has been shown to be regulated by MT stability, as Kif5B (kinesin 1) normally transports cargo selectively into axons, however after treatment with paclitaxel also targets dendrites (Kapitein et al., 2010; Konishi & Setou, 2009). Kif5B has also been connected to lysosome transport in autophagy, and depletion of Kif5B led to perinuclear accumulation of autophagosomes in cancer cells (Cardoso et al., 2009; Guardia et al., 2016). In the context of insulin vesicle transport a connection between Kif5B and GABARAP has been proposed, as GABARAP appears to promote vesicle trafficking by Kif5B (Asano et al., 2014). Considering our results, it is conceivable that GABARAP not only presents transport vesicles to KIF5B by connecting vesicles and MTs and thereby stabilizing the kinesin-cargo complex, but that high local GABARAP concentrations additionally stabilize MTs and thereby support Kif5B binding and corresponding anterograde vesicular transport.

Considering that the members of the LC3 subfamily of ATG8 proteins have been first described as light chains of the MT interacting proteins MAP1A and MAP1B (Kouno et al., 2005; Mann & Hammarback, 1994), further connections between ATG8s and MTs are not far-fetched. Accordingly, the connection between autophagy and the cytoskeleton has been extensively studied and reviewed (Kast & Dominguez, 2017; Mackeh et al., 2013; Monastyrska et al., 2009). Regarding the autophagy-unrelated functions, an interplay between GABARAP and MTs has been suggested for the anterograde transport of the GABA_A_, angiotensin II type 1 and κ opioid receptors (Chen et al., 2011; Cook et al., 2008; Leil et al., 2004). The LC3B-mediated transport of melanosomes along MTs and detachment from MTs by ATG4B-mediated delipidation is another example of this connection (Ramkumar et al., 2017).

In summary, while conventionally tagged GABARAP, e.g. *B*-G, does not visually associate with MTs (Figure 5C-I), possibly because the adverse effect of the tag is too severe to be compensated by the following single GABARAP, G*-B*-G accumulates on MTs (Figure 5C-II), presumably owing to the bivalent nature of the construct. This is supported by the fact that *B*-G-G shows comparable MT association. Drechsler et al (2019) recently reported that multivalence is a critical property of MAPs, conferring MT-bundling abilities, optionally by bridging individual MTs (Figure 5C-III). In the case of untagged, endogenous GABARAP, we hypothesize that the soluble form associates with MTs through charge-mediated enrichment, albeit with relatively low affinity (Figure 5C-IV). GABARAP-decorated transport vesicles, however, could present multiple GABARAPs in proximity to MTs, resulting in significant avidity through multivalence (Figure 5C-V). Such multiple GABARAP-MT interactions might therefore assist in keeping transport vesicles on the MT track, possibly promoting transport (Fig. 5C-VI).

As both impaired MT dynamics and defective autophagy as well as their interplay have been linked to human disease (Arduino et al., 2012; Guo et al., 2018; Naren et al., 2023), understanding how GABARAP and other ATG8 proteins influence MT stability and related processes may also provide novel insights into pathophysiology and suggest improved strategies of intervention.

As exemplified with G-*B*-G, the split tandem strategy offers a promising tool to investigate putative connections of the entire range of ATG8 family proteins with MTs, and the field of application is certainly open towards areas beyond the cytoskeleton.

## Data and material availability

Original imaging data is available on BioImage Archive at https://www.ebi.ac.uk/biostudies/bioimages/studies/S-BIAD952. All described plasmids are deposited at Addgene.

## Acknowledgment

We thank Tom Boissonnet and Vanessa Fuchs (Center for Advanced Imaging (CAi), Heinrich Heine University Düsseldorf, Germany) for their kind assistance in using OMERO and in submitting the data to the BioImage Archive.

## Author Contributions

Conceptualization: S.H., O.H.W., A.Ü., T.G.; validation: S.H., O.H.W., A.Ü.; formal analysis: A.Ü.; investigation: A.Ü., L.G., O.H.W.; resources: D.W.; data curation: A.Ü.; writing— original draft preparation: A.Ü., S.H.; writing – review and editing: S.H., O.H.W., A.Ü, T.G., L.G., D.W.; visualization: A.Ü.; supervision: S.H., O.H.W.; project administration: D.W.; funding acquisition: D.W.

ATG8: autophagy related protein 8
FP: fluorescent protein
GABARAP: γ-aminobutyric acid type A receptor-associated protein
HA-tag: hemagglutinin tag
Icorr: Index of correlation
KO: knockout
LDS: LIR docking site
LIR: LC3-interacting region
MAP1LC3/LC3: microtubule associated protein 1 light chain 3
MAP: microtubule associated protein
MT: microtubule
nMDP: normalized mean deviation product
ROI: region of interest
SiR: Silicon rhodamine

