## Supplementary Information for "Highlighting the hidden: A tagging strategy for monitoring the association of GABARAP with microtubules in living cells"

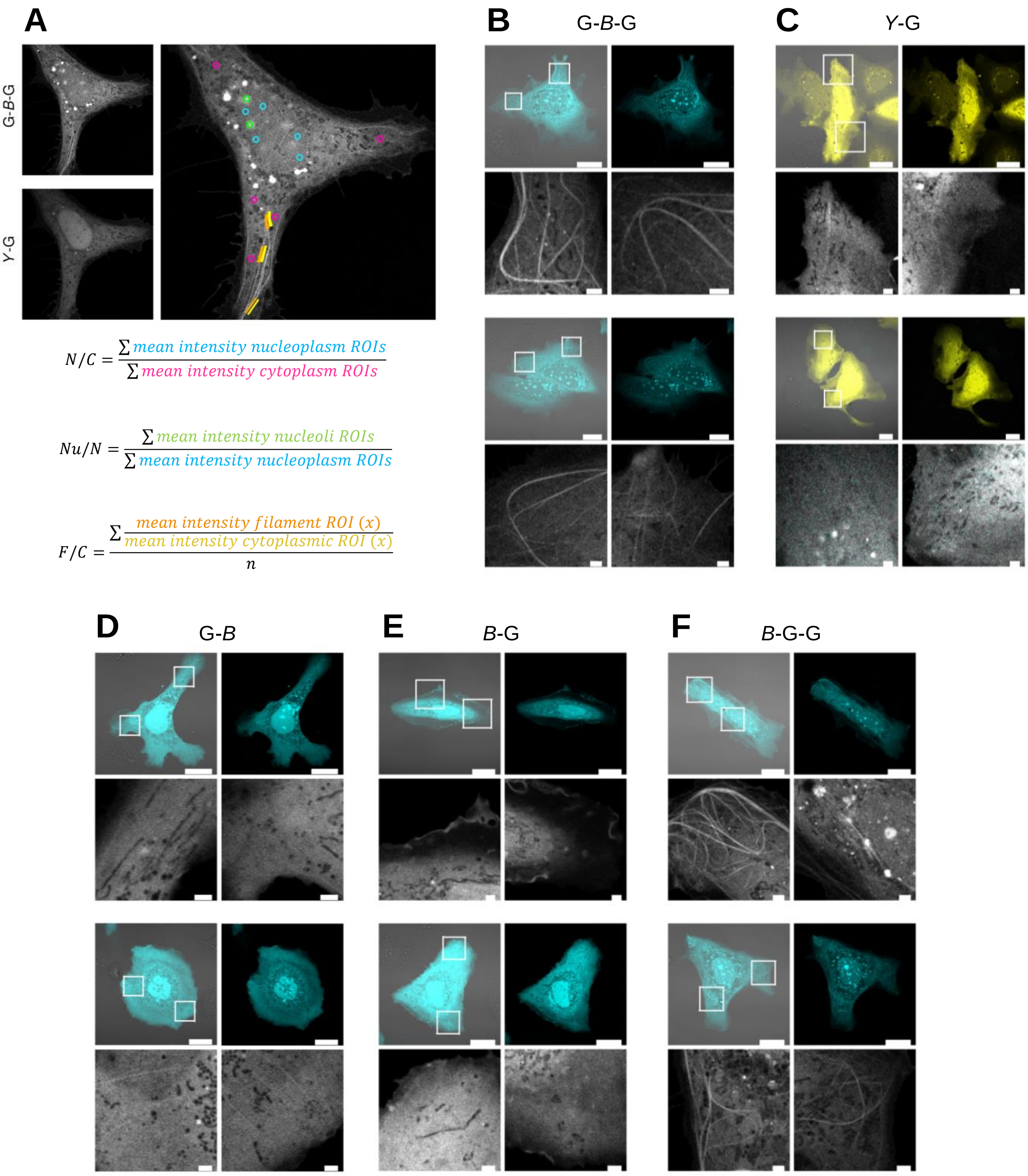


**Figure S1.** (**A**) Example of cytoplasmic, nucleoplasmic, nucleoli and filament ROIs in a Huh$^{KO}$ cell expressing G-*B*-G and *Y*-G as well as the formulas used for calculation of N/C, Nu/N and F/C ratios. All images and ROIs of cells analyzed accordingly can be found on BioImageArchive. (**B-F**) Live cell images of Huh$^{KO}$ cells expressing G-*B*-G (**B**), *Y*-G (**C**), *B*-G (**D**), G-*B* (**E**) *B*-G-G (**F**). Whole cells (merge with transmitted and G-*B*-G channel) as well as detailed ROIs are shown for two cells per construct. Scale bars represent 20 µm for whole cells and 2 µm for zoom ins.

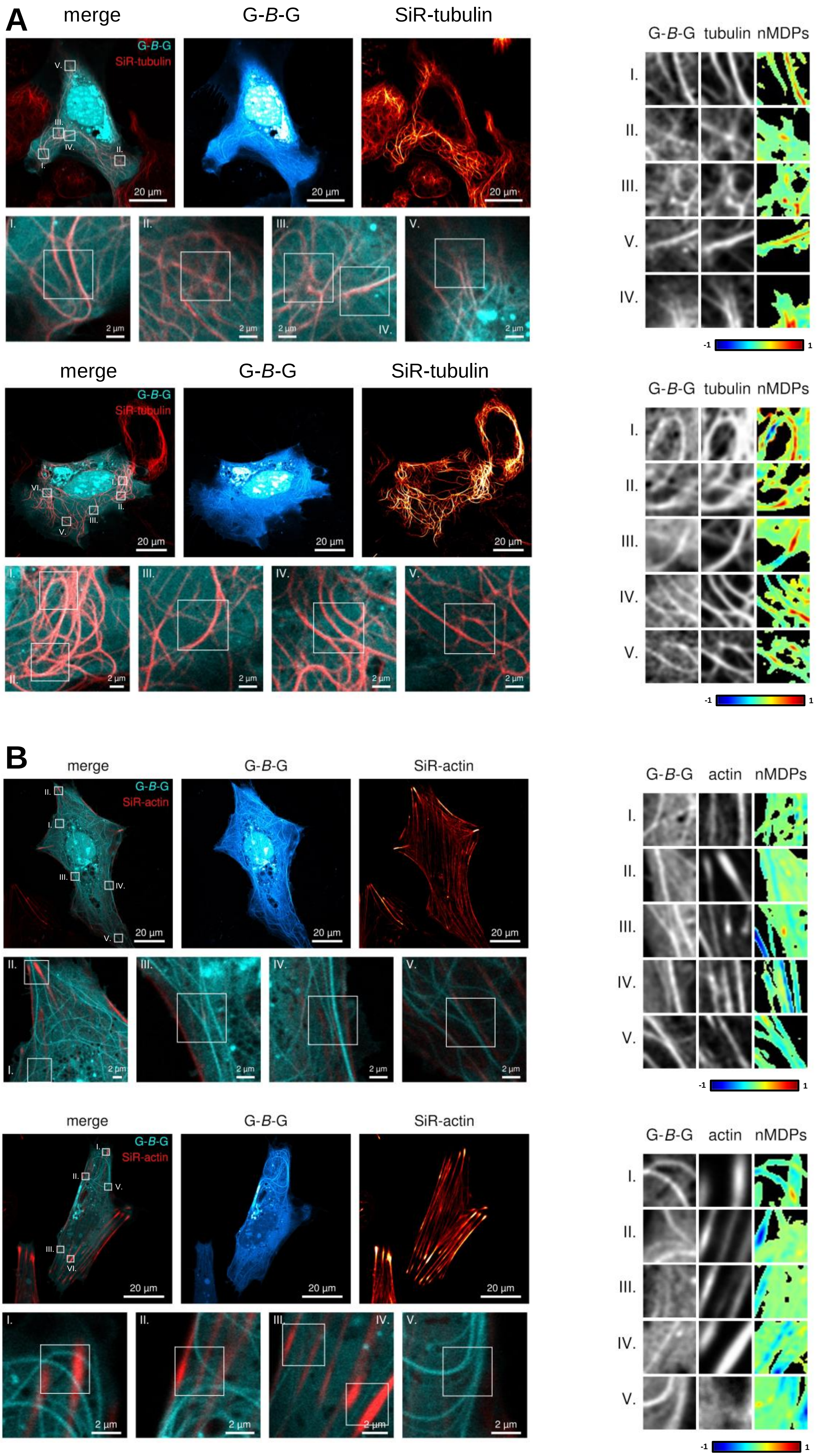


**Figure S2.** Continued on the following page.

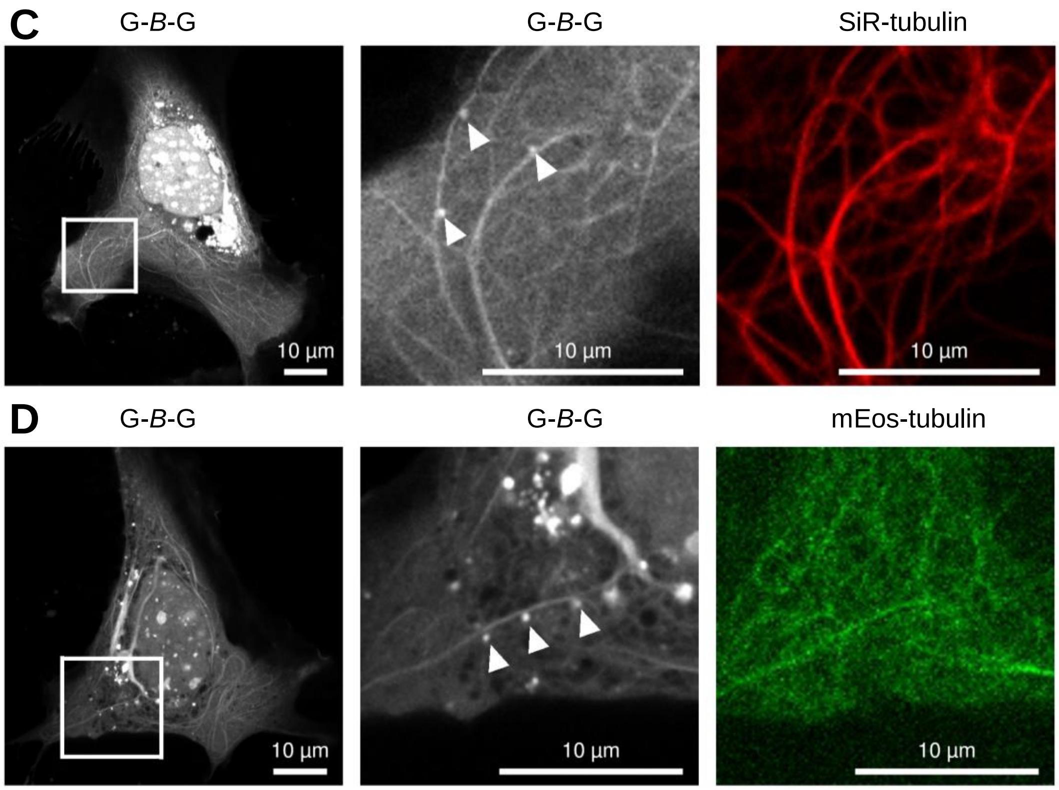


**Figure S2.** Live cell images of Huh$^{KO}$ cells expressing G-*B*-G and stained either with SiR-tubulin (**A**) or SiR-actin (**B**) together with their colocalization colormaps. The corresponding Icorr values are included within the graph given in Fig. 2D. (**C-D**) Live cell images of Huh$^{KO}$ cells expressing G-*B*-G and either co-stained with SiR-tubulin (**C**) or co-transfected with a plasmid encoding mEos-tubulin (**D**) Zoom in images show microtubules decorated with G-*B*-G positive puncta, likely transport vesicles.

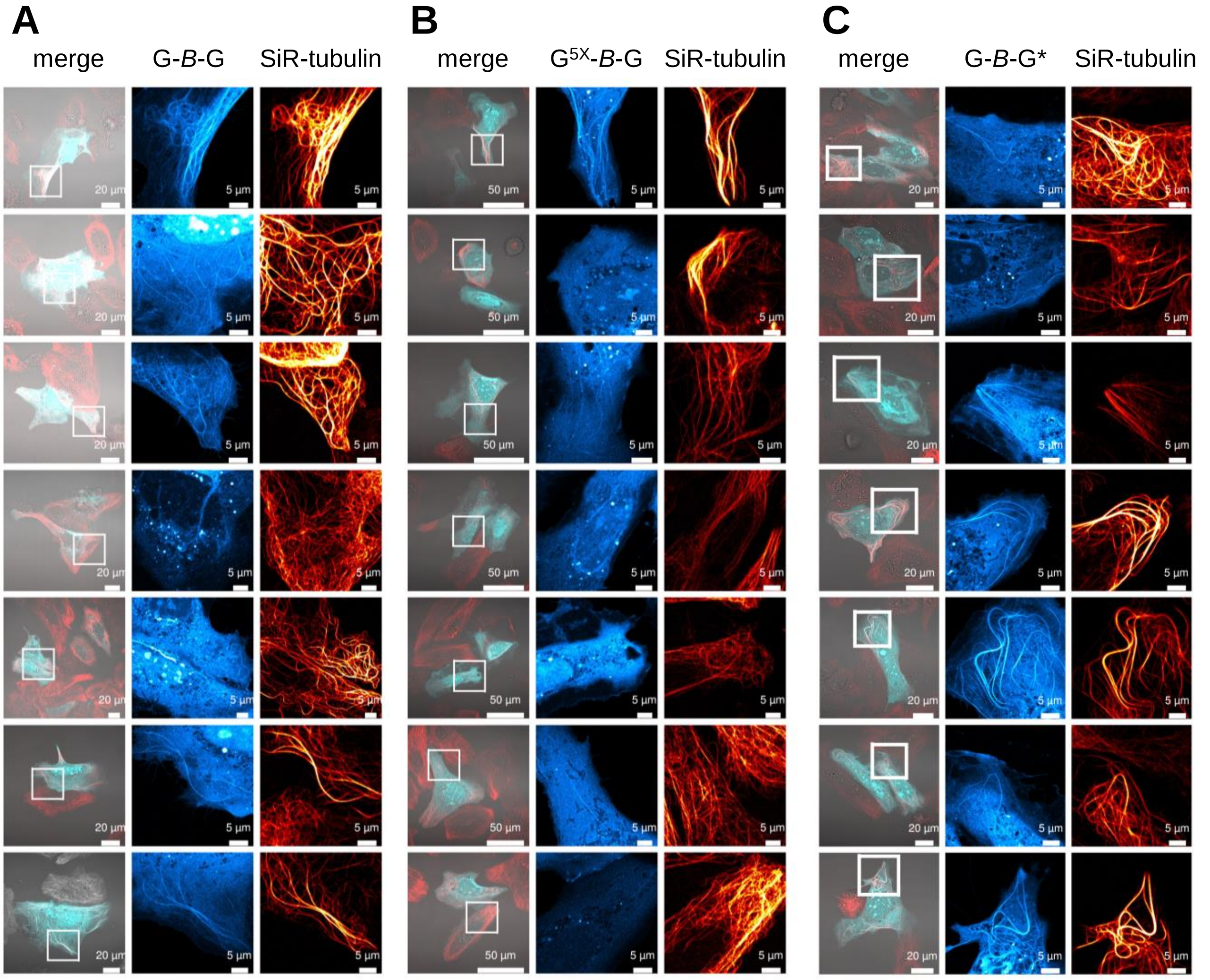


**Figure S3.** Continued on the following page.

D

E

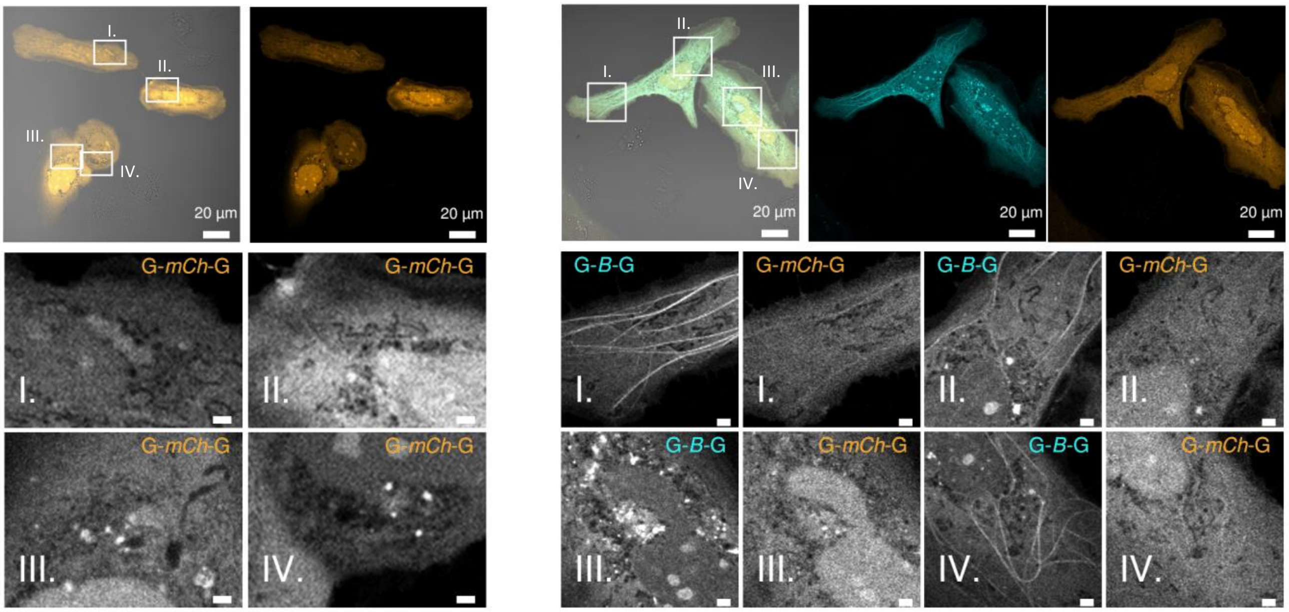


F

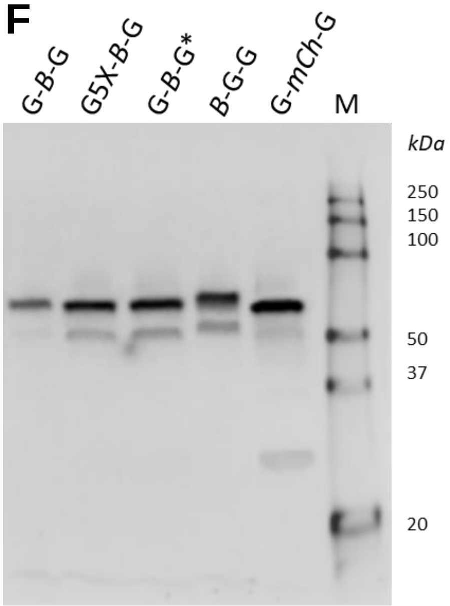


**Figure S3.** (**A-C**) Live cell images of Huh$^{KO}$ cells expressing G-*B*-G (**A**), G$^{5X}$-*B*-G (**B**) or G-*B*-G* (**C**) and co-stained with SiR-tubulin. All presented cells were used for determination of F/C ratios shown in Figure 3D. Corresponding ROIs for quantification can be accessed on BioImageArchive. (**D-E**) Huh$^{KO}$ cells expressing split tandem construct (**D**) G-*mCh*-G alone and (**E**) together with G-*B*-G. Scale bars in zoom ins: 2 µm (**F**) Full length expression of different split tandem constructs (G-*B*-G, G5X-*B*-G, G-*B*-G*, *B*-G-G and G-*mCh*-G) was confirmed by Immunoblot from whole cell lysates of Huh$^{KO}$ cells transfected with the respective plasmids and and staining with a monoclonal GABARAP antibody. The most intense bands were detected at the expected size of the construct at approximately 55 kDa. The signal at about 25 kDa is unlikely to contribute to fluorescence signal in cells.

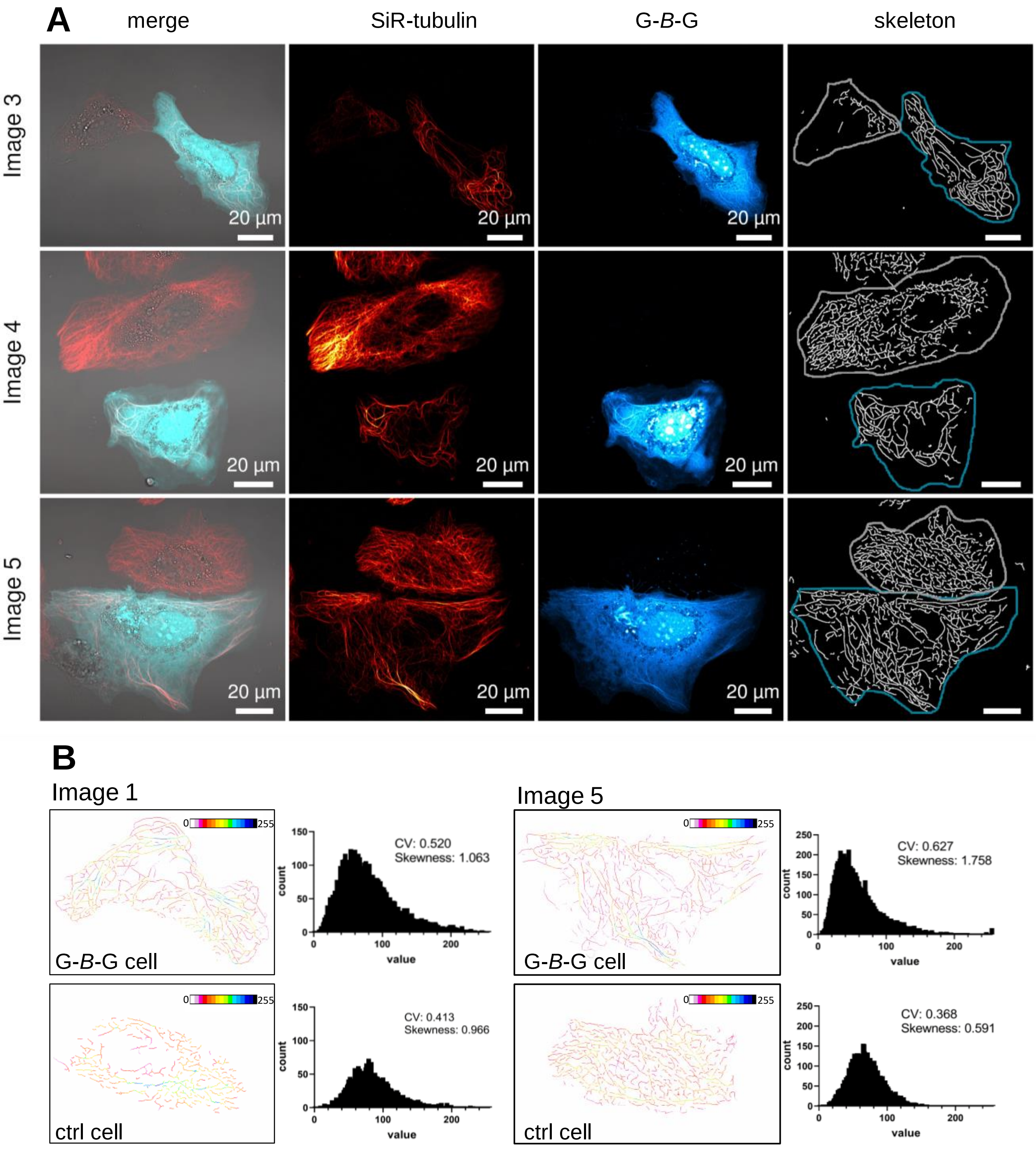


**Figure S4.** Continued on the following page.

C

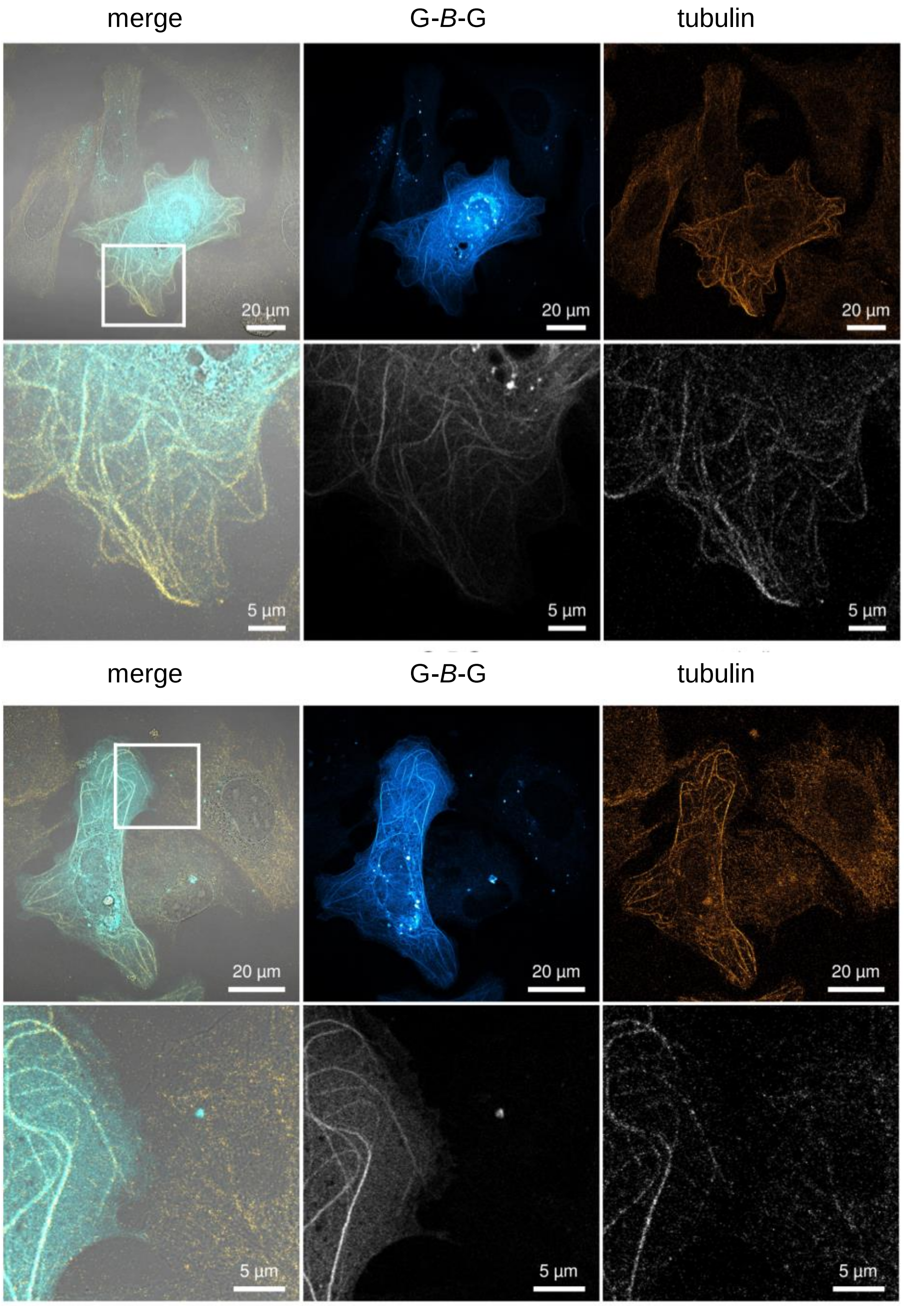


**Figure S4.** (**A**) Images of Huh$^{KO}$ cell pairs, one expressing G-*B*-G and a second untransfected control cell, stained with SiR-tubulin. Skeletons, thresholded and dilated for visualization, for analysis of branch length as shown in Figure 4A are displayed. (**B**) For image 1 and 5, color coded skeletons according to tubulin intensity values and corresponding cytoskeleton bundling parameters are shown analogous to Figure 4E-G. (**C**) Images of fixed Huh cells transiently transfected with plasmid encoding G-*B*-G and stained with an antibody against tubulin. Cells showing strong G-*B*-G expression present an altered tubulin staining pattern.

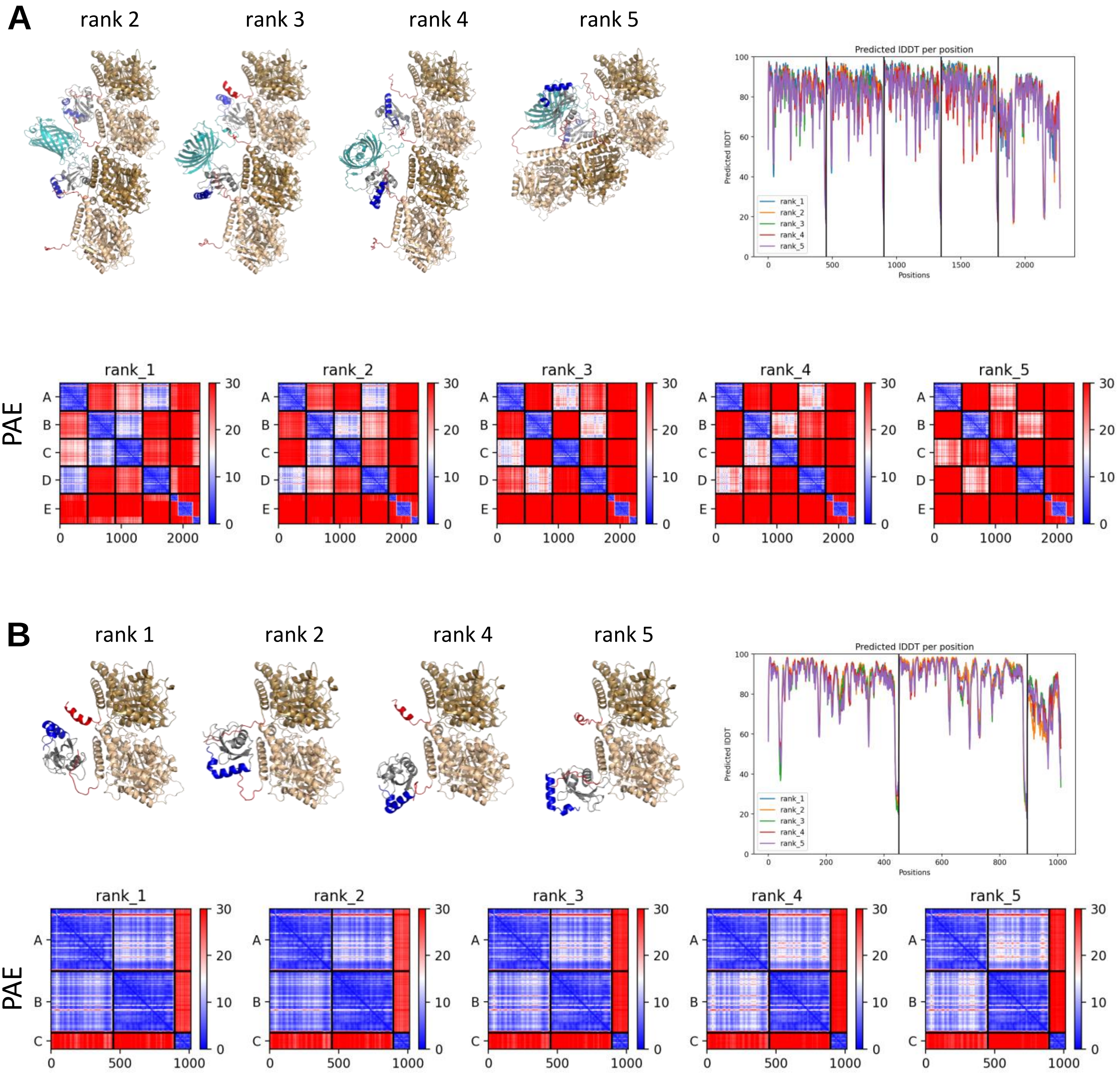


**Figure S5.** Continued on the following page.

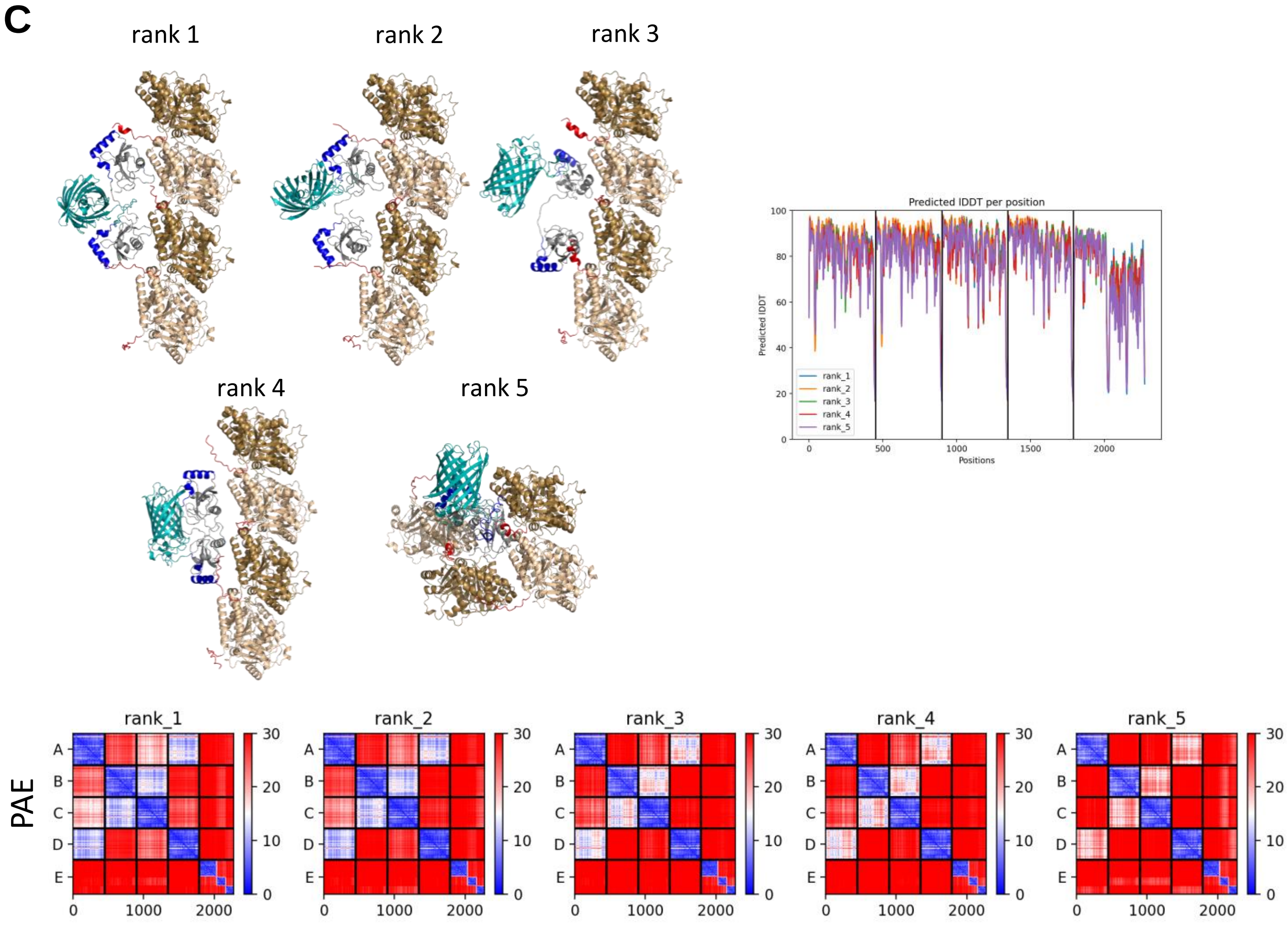


**Figure S5.** ColabFold data not shown in the main manuscript, including additional models as well as per-residue confidence metrics (predicted local difference distance test, lDDT, and predicted aligned error, PAE). (**A**) G-*B*-G with a TBA1A/TBB5 tetramer; rank-1 model in Fig. 5A. (**B**) GABARAP with a TBA1A/TBB5 dimer; rank-3 model in Fig. 5B. (**C**) B-G-G with a TBA1A/TBB5 tetramer.

### SI Methods

#### Immunocytochemistry

Transfected Huh-7.5 GABARAP SKO cells were fixed for 10 min with precooled methanol and subsequently for 1 minute with precooled acetone, both at -20°C. After two washes with PBS (137 mM NaCl, 2.7 mM KCl, 1.8 mM $KH_2PO_4$, 10 mM $Na_2HPO_4$, pH 7.4), non-specific binding sites were blocked with 5% (w/v) BSA in PBS for 1 h. Cells were incubated with primary antibodies (anti-GABARAP rabbit monoclonal [13733, Cell Signaling Technology], anti-GABARAP rabbit polyclonal [18723-I-AP, Proteintech], anti-β-tubulin-CY3 mouse monoclonal [C4585, Sigma Aldrich]) diluted 1:100 in PBS containing 1% (w/v) BSA (A1391, AppliChem) and 0.3% (v/v) Triton X-100 (A4975, AppliChem) for 2 h or overnight and afterwards washed twice with PBS. If required, cells were stained with secondary antibody (anti-rabbit-Alexa647 [ab150083, abcam]) diluted 1:200 in the abovementioned solution for 1 h. Following two washes with PBS, cells were imaged and stored in PBS containing 0.05% (w/v) sodium azide.

#### Immunoblotting

Transfected Huh-7.5 GABARAP SKO cells were harvested by trypsination, washed once in PBS and lysed with NP40 buffer (20 mM Tris HCl, 200 mM NaCl, 1 mM EDTA, 0.5% NP40, 1 mM PMSF and 1x Halt Protease and Phosphatase inhibitor [78442, Thermo Fisher Scientific]) by incubation on ice for 30 min and vigorous pipetting every 10 min. Unsoluble cell part were sedimented by centrifugation for 10 min at 17,000 x g. Total protein concentration of supernatant was determined by BCA assay (23225, Thermo Scientific). Samples were separated by SDS-PAGE using precast stain-free gels (4568124, BioRad), transferred to 0.2 µM PVDF membranes (1704156, BioRad) and blocked in 5% BSA (A1391, AppliChem) in TBS-T (136 mM NaCl, 2.7 mM KCl, 24.7 mM Tris-HCl, pH 7.4, 0.05% Tween-20 [A4974, Applichem]) for 1 h. Membrane was incubated with primary anti GABARAP antibody (13733, Cell Signaling Technology) overnight at 4°C, washed 3 times with TBS-T, incubated with secondary antibody (goat anti-rabbit, HRP-conjugated [P0448, Dako]) for 1 h at room temperature and again washed thrice with TBS-T. Signals were visualized with Clarity western ECL substrate (1705061, BioRad).
